# A traditional medicine, Respiratory Detox Shot (RDS), inhibits the infection of SARS-CoV, SARS-CoV-2, and the Influenza A virus *in vitro*

**DOI:** 10.1101/2020.12.10.420489

**Authors:** Brian Hetrick, Dongyang Yu, Adeyemi A. Olanrewaju, Linda D. Chilin, Sijia He, Deemah Debbagh, Yuan-Chun Ma, Lewis A. Hofmann, Ramin M. Hakami, Yuntao Wu

**Affiliations:** National Center for Biodefense and Infectious Diseases, School of Systems Biology, George Mason University, Manassas, VA 20110, USA; Virongy LLC, Manassas, VA 20109, USA; Dr. Ma’s Laboratories Inc. Burnaby, BC V5J 0E5, Canada; World Health Science Organization, Leesburg, VA 20176, USA

## Abstract

The ongoing global pandemic of coronavirus disease 2019 (COVID-19) has resulted in the infection of over 60 million people and has caused over 1.4 million deaths as of December 2020 in more than 220 countries and territories. Currently, there is no effective treatment for COVID-19 to reduce mortality. We investigated the potential anti-coronavirus activities from an oral liquid of traditional medicine, Respiratory Detox Shot (RDS), which contains mostly herbal ingredients traditionally used to manage lung diseases. Here we report that RDS inhibited the infection of target cells by SARS-CoV and SARS-CoV-2 pseudoviruses, and by infectious wild-type SARS-CoV-2. We further demonstrated that RDS inhibits viral early infection steps. In addition, we found that RDS can also block the infection of target cells by Influenza A virus. These results suggest that RDS may broadly inhibit the infection of respiratory viruses.

## INTRODUCTION

The ongoing coronavirus disease 2019 (COVID-19) global pandemic has afflicted more than 60 million people in over 220 countries and territories, resulting in more than 1.4 million deaths as of December 2020. Currently, there is no effective treatment for COVID-19 to reduce mortality. The newly emerged viral pathogen causing COVID-19 is the coronavirus SARS-CoV-2 (*1*), a sister virus of SARS-CoV in the species of *Severe acute respiratory syndrome-related coronavirus* (*2, 3*). Both SARS-CoV and SARS-CoV-2 emerged in China; SARS-CoV was first identified in Guangdong Province in November 2002 (*4-6*), and SARS-CoV-2 was first identified in Wuhan in December 2019 (*1, 7, 8*). In both coronavirus-caused pandemics, traditional Chinese medicines (TCM) have been widely used in China for the urgent management of coronavirus diseases. For the current COVID-19 pandemic, greater than 85% of SARS-CoV-2 infected patients in China have received TCM treatments of some sort (*9, 10*). Whether many of the TCMs used have active anti-coronavirus properties and are clinically effective are important questions that have not been fully answered. Lack of systemic studies, both *in vitro* and *in vivo*, have hampered the development and rational use of TCMs as effective therapeutics for the treatment of coronavirus diseases.

To identify potential anti-SARS-CoV-2 activities from traditional herbal medicines, we screened multiple herbal extracts, and discovered anti-SARS-CoV and anti-SARS–CoV-2 activities from an oral liquid, Respiratory Detox Shot (RDS), a commercial food supplement in the United States. RDS is used to manage the general wellness of the human respiratory system, and contains multiple herbal ingredients, such as *Panax ginseng and Schizonepeta tenuifolia*, that are Chinese herbal medicines traditionally used to manage inflammation and lung diseases (*11-13*). Here we report that RDS inhibited the infection of target cells by SARS-CoV and SARS-CoV-2 pseudoviruses, and by infectious wild-type SARS-CoV-2. We further demonstrate that RDS inhibits viral early infection steps likely by directly inactivating virion or blocking viral entry. In addition, we found that RDS potently blocks the infection of Influenza A virus. These results suggest that RDS may broadly inhibit the infection of respiratory viruses.

## RESULTS

To discover potential anti-SARS-CoV-2 activities from traditional herbal medicines, we screened extracts from approximately 40 medicinal herbs, using a SARS-CoV-2 S protein pseudotyped lentivirus (*14, 15*) and human lung A549(ACE2) target cells, in which the human ACE2 gene is over-expressed through lentiviral vector-mediated stable transduction. The lentipseudoviruses use either the green fluorescent protein (GFP) or luciferase (Luc) as the reporter, and were validated with a broad-spectrum antiviral entry inhibitor, Arbidol (*16*), and human antiserum against SARS-CoV-2 (**Fig. 1A** and **1C**). We were able to detect inhibition of SARS-CoV-2 pseudovirus by Arbidol and the antiserum, while we did not find inhibition from any of the herbal extracts tested even in the presence of high toxicity from some of them (**Fig 1A to 1C**).

**Figure 1.**
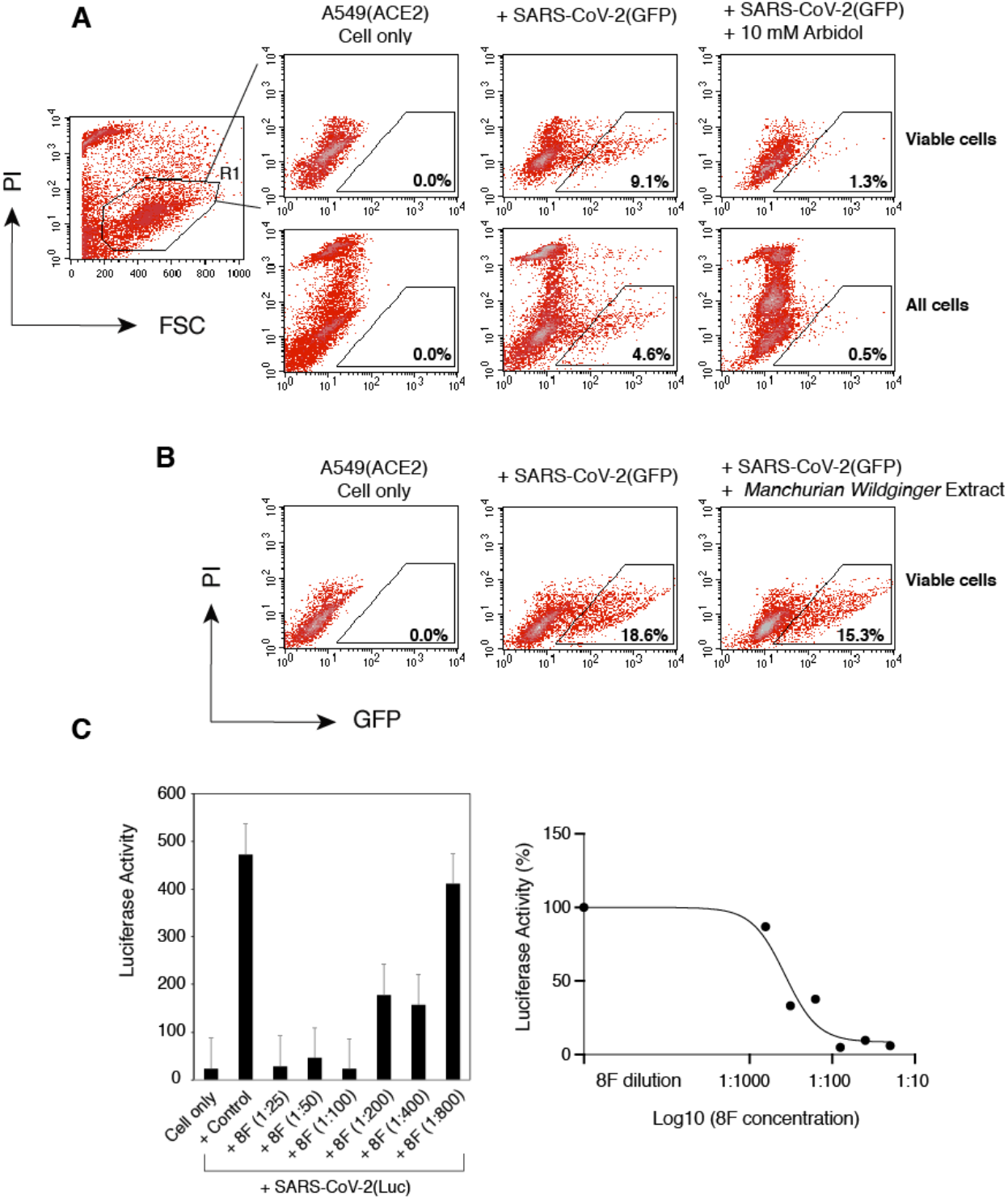
Validation of SARS-CoV-2 S protein pseudotyped reporter viruses for the screening and quantification of antiviral drugs and neutralization antibodies. (**A**) A lentiviral particle, SARS-CoV-2(GFP) that was pseudotyped with the SARS-CoV-2 S protein, was used to infect A549(ACE2) target cells. GFP was used as the reporter to quantify viral infection, and was measured at 48 hours post infection with flow cytometry. Propidium iodide (PI) was added during flow cytometry to stain for dying and dead cells. Arbidol (10 mM) was tested in the system for blocking viral infection. GFP+ cells were quantified only in the viable cell population or in the whole cell population. (**B**) An example of using SARS-CoV-2(GFP) to screen for TCMs. Extract from *Manchurian Wildginger* (2 mg/ml) was used to pretreat cells which were infected in the presence of *M. Wildginger*. After infection, cells were cultured in the absence of *M. Wildginger* for 48 hours. GFP expression was quantified. (**C**) A lentiviral particle, SARS-CoV-2(Luc) pseudotyped with the SARS-CoV-2 S protein, was used to infect A549(ACE2) target cells. Luciferease (Luc) was used as the reporter to quantify viral infection. Luc was measured at 72 hours post infection. The SARS-CoV-2 neutralizing antiserum 8F was serially diluted and incubated with viral particles for 1 hour. The complex was added to infect cells. Luc expression was quantified at 72 hours post infection.

We further screened possible anti-SARS-CoV-2 activity from an oral liquid of a traditional medicine, Respiratory Detox Shot (RDS), which contains nine ingredients (*Lonicera japonica, Forsythia suspensa, Panax ginseng, Schizonepeta tenuifolia, Scrophularia ningpoensis, Prunus armeniaca, Polistes mandarinus saussure, Gleditsia sinensis, Glycyrrhiza uralensis*) traditionally used in China to manage lung diseases (*11-13*). A549(ACE2) cells were pretreated with serially diluted RDS, and then infected in the presence of RDS for 4-6 hours. Following infection, cells were cultured in the absence of RDS, and then quantified for the inhibition of viral infection by flow cytometry at 48 and 72 hours. To control for cytotoxicity, propidium iodide (PI) was used to stain for dying and dead cells, and GFP+ cells were analyzed only in the viable cell population. As shown in **Fig. 2**, we observed RDS dosage-dependent inhibition of the SARS-CoV-2(GFP) pseudovirus. To confirm these results, we repeated the infection using Vero E6 cells that endogenously express ACE2; Vero E6 supports productive SARS-CoV and SARS-CoV-2 infection and is commonly used in studies of coronaviruses (*7*). Given the low infectivity of pseudoviruses for Vero E6 in the absence of ACE2 overexpression (*15, 17, 18*), we also used a Luc reporter pseudovirus, in which the reporter expression is driven by HIV-1 LTR and Tat for higher reporter sensitivity and signal to noise ratio. As shown in **Fig. 3A**, using the Luc reporter pseudovirus and Vero E6, we observed RDS dosage-dependent inhibition of infection, and the IC50 (50% inhibition dosage) was determined to be at 1: 230 RDS dilution (**Fig. 3B**). We also quantified effects of RDS on Vero E6 cell viability, and the LC50 (50% cell death dosage) was determined to be at 1: 11.8 RDS dilution (**Fig. 3C**). To further validate the results obtained from using pseudoviruses, we tested the ability of RDS to block the infection of wild-type SARS-CoV-2. As shown in **Fig. 3D**, RDS also blocked the infection of Vero E6 cells by SARS-Cov-2. RDS greatly diminished the formation of viral plaques at dosages above 1:40 dilution, and reduced viral replication approxmiately 2-3 logs at 1:40 dilution. Together, the results from SARS-CoV-2 pseudoviruses and the wild-type virus demonstrated that RDS contains active ingredients inhibting SARS-CoV-2 infection, likely by directly inactivating virons or by blocking viral early infection steps.

**Figure 2.**
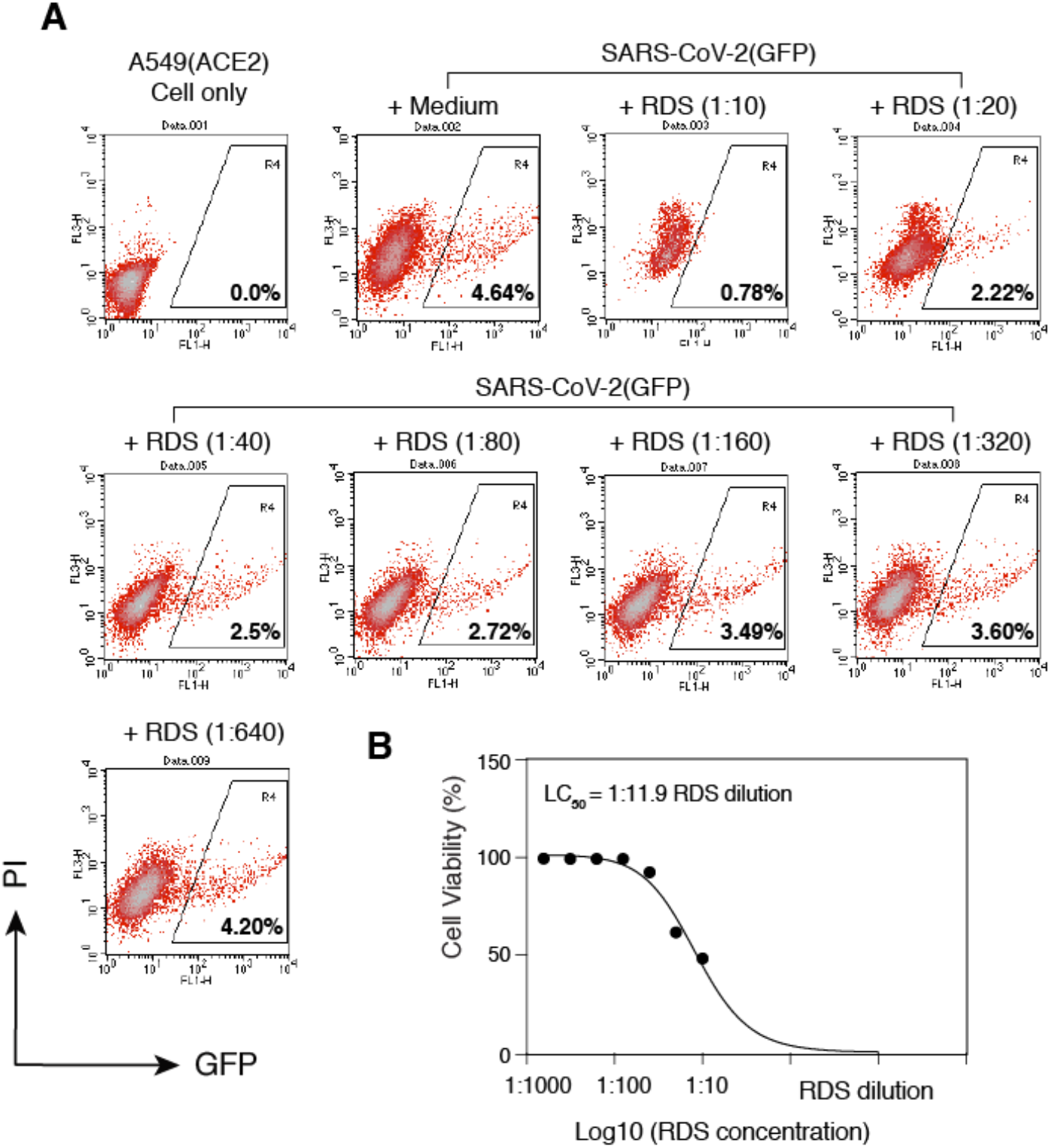
RDS inhibits SARS-CoV-2(GFP) pseudovirus infection of A549(ACE2) cells. (**A**) A549(ACE2) cells were treated with serially diluted RDS for 30 minutes, and then infected with SARS-CoV-2(GFP) pseudovirus. Cells were washed to remove the virus and RDS, and cultured in the absence of RDS. Inhibition of viral infection was quantified by flow cytometry. Uninfected cell and SARS-CoV-2(GFP)-infected but RDS-untreated cells were used as controls. The percentages of GFP+ cells are shown. PI, propidium iodide. (**B**) Quantification of the cytotoxicity of RDS. A549(ACE2) cells were treated with serially diluted RDS for 4 hours, washed to remove RDS, and cultured in the absence of RDS for 48 hours. Cells were stained with propidium iodide to identify dying and dead cells, and analyzed with flow cytometry. The dose-response cytotoxicity curve was plotted, and the LC50 of RDS was calculated to be at 1:11.9 dilution.

**Figure 3.**
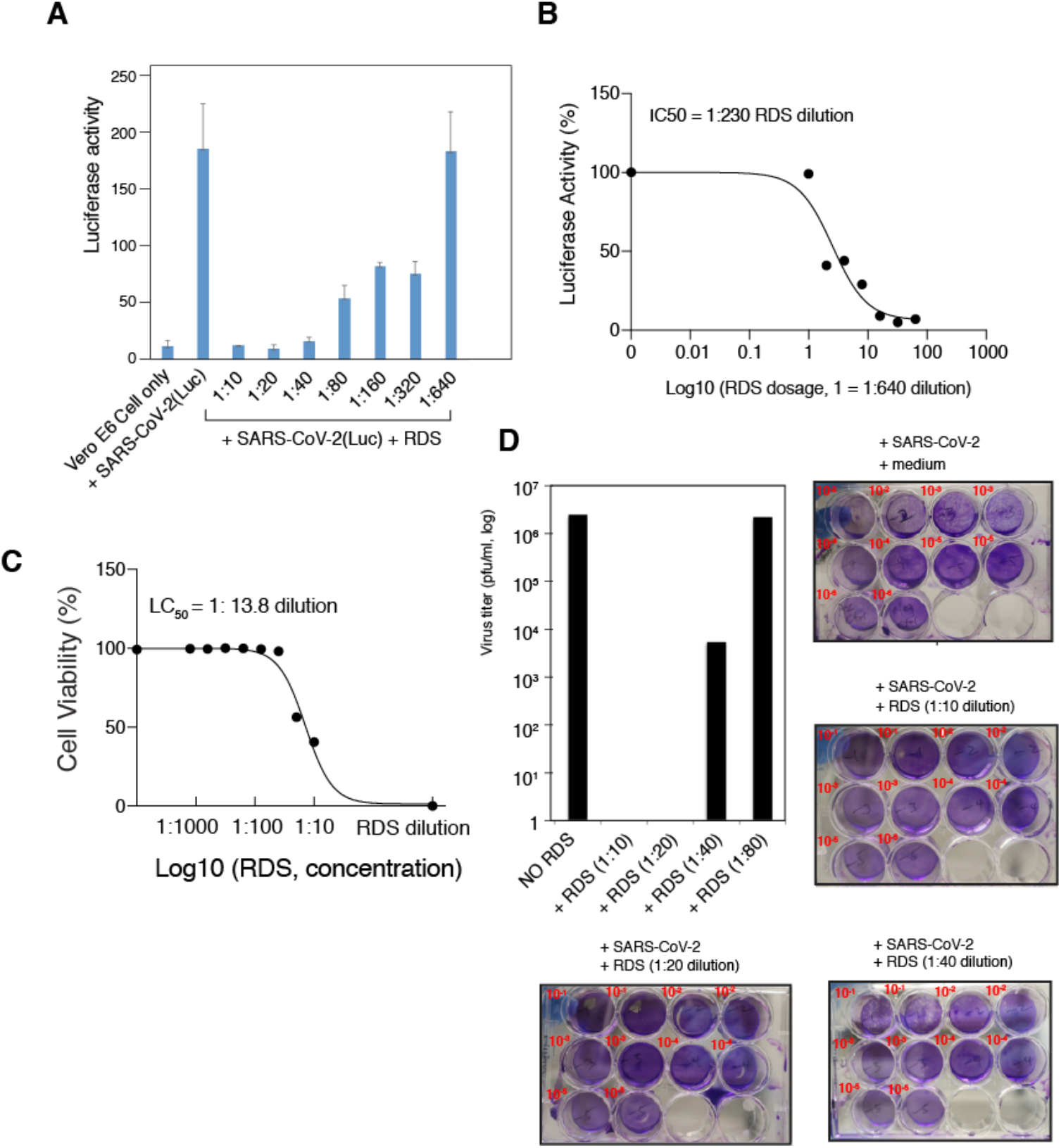
RDS dosage-dependent inhibition of SARS-CoV-2(Luc) pseudovirus and wild-type SARS-CoV-2. (**A** and **B**) Vero E6 cells were pretreated with serially diluted RDS, and infected with SARS-CoV-2(Luc) pseudovirus. Cells were washed to remove the virus and RDS, and cultured in the absence of RDS. Inhibition of viral infection was quantified at 72 hours post infection by luciferase assay. Uninfected cell and SARS-CoV-2-Luc-infected but RDS-untreated cells were used as controls. The assay was performed in triplicate. The dose-response curve was plotted, and the IC50 of RDS was quantified to be at 1:230 dilution. (**C**) The cytotoxicity of RDS on Vero E6 cells was also quantified using propidium iodide staining and flow cytometry. Cells were treated with serially diluted RDS for 4 hours, washed to remove RDS, and cultured in the absence of RDS for 72 hours. The dose-response cytotoxicity curve was plotted, and the LC50 of RDS was calculated to be at 1:13.8 dilution. (**D**) RDS inhibits wild-type SARS-CoV-2 infection. Vero E6 cells were pretreated with serially diluted RDS, and infected with SARS-CoV-2 in the presence of RDS. Inhibition of viral replication was quantified by plaque assays of the virus released at 48 hours post infection.

We also tested the ability of RDS to block the infection of SARS-CoV, using a GFP reporter lentivirus pseudotyped with the SARS-CoV spike protein (*15*). Human A549(ACE2) cells was used as the target cells, which were pretreated with serially diluted RDS, and then infected with SARS-CoV(GFP) reporter pseudovirus for 4-6 hours. Following infection, cells were cultured in the absence of RDS, and then quantified for the inhibition of viral infection by flow cytometry. Similarly, propidium iodide was used to exclude dying and dead cells, and GFP+ cells were analyzed only in the viable cell population. As shown in **Fig. 4A**, we observed RDS dosage-dependent inhibition of SARS-CoV(GFP) pseudovirus. We further confirmed these results and quantified the RDS-mediated inhibition with a Luc reporter SAS-CoV pseudovirus, SARS-CoV(Luc). We observed dosage-dependent RDS inhibition of SARS-CoV(Luc), and the IC50 was determined to be at 1:70.88 RDS dilution (**Fig. 4B and 4C**).

**Figure 4.**
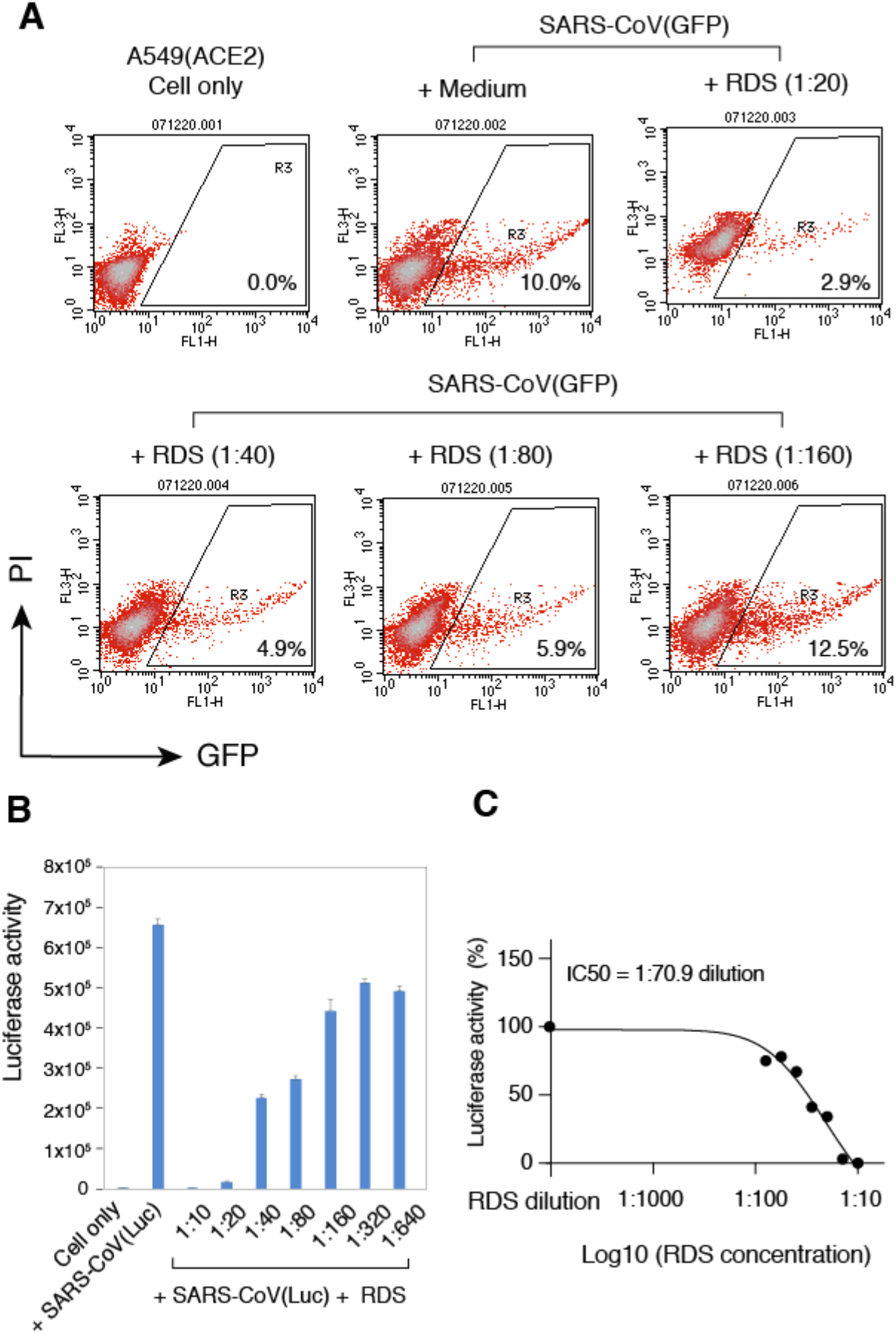
RDS inhibits SARS-CoV pseudovirus infection of A549(ACE2) cells. (**A** and **B**) Cells were pretreated with serially diluted RDS, and infected with SARS-CoV(GFP) (**A**) or SARS-CoV(Luc) (**B**) pseudovirus. Cells were washed to remove the virus and RDS, and cultured in the absence of RDS. Inhibition of viral infection was quantified at 48 hours or 72 hours post infection by flow cytometry or luciferase assay. The assay was performed in triplicate. The dose-response curve was plotted, and the IC50 of RDS was determined to be at 1:70.9 dilution (**C**).

Given that both SARS-CoV and SARS-CoV-2 use ACE2 to infect target cells, we also tested whether the anti-viral activity of RDS is specific to coronaviruses interacting with ACE2. For this purpose, we tested an unrelated, negative-sense RNA virus, Influenza A, which uses viral hemagglutinin (HA) and cellular α-sialic acid to infect target cells. To assemble influenza A virus, eight vectors expressing each of the segments of the influenza A/WSN/33 (H1N1) genome plus a GFP-reporter vector were cotransfected into HEK293T cells. Viral particles were harvested and used to infect target MDCK cells in the presence of RDS. As shown in **Fig. 5A**, we observed dosage-dependent inhibition of Influenza A virus by RDS. RDS completely blocked viral infection at dilutions of 1:40 and 1:80, and partially inhibited Influenza A at 1:160. The cytotoxicity LC50 of RDS on MDCK cells was determined to be at 1:18.5 dilution (**Fig. 5B**). These results suggest that the anti-viral activities of RDS are not virus-specific, and may broadly inhibit multiple respiratory viruses such as coronaviruses and Influenza A.

**Figure 5.**
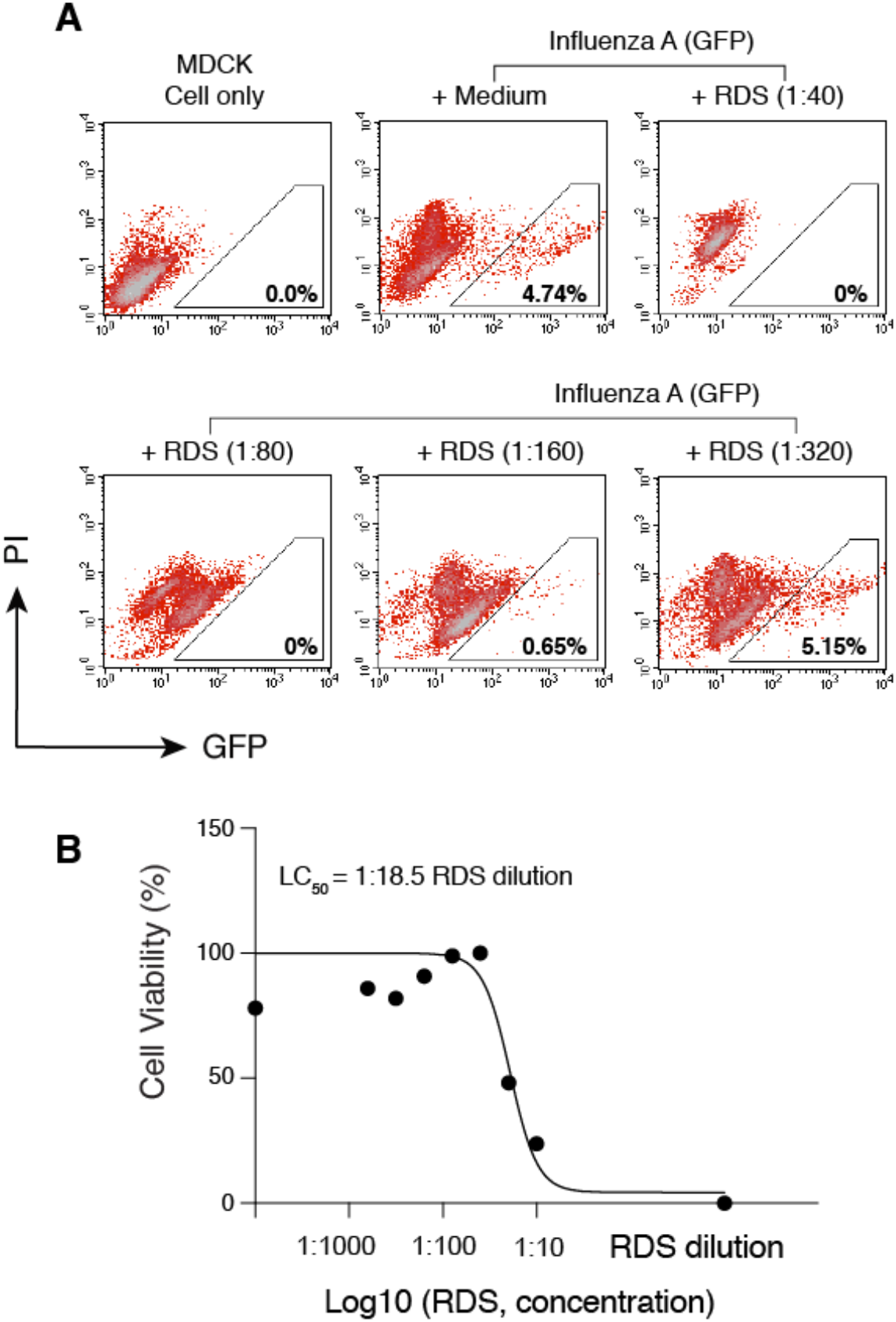
RDS inhibits influenza A virus infection of MDCK cells. (**A**) MDCK cells were pre-treated with serially diluted RDS for 30 minutes, and then infected with FluA(GFP) virus. Following infection, cells were cultured in the presence of RDS. Inhibition of viral infection was quantified at 36 hours post infection with flow cytometry. Uninfected cell and FluA(GFP)-infected but RDS-untreated cells were used as controls. The percentages of GFP+ cells are shown. PI, propidium iodide. (**B**) The cytotoxicity of RDS on MDCK cells was also quantified using MTT assay. The dose-response cytotoxicity curve was plotted, and the LC50 of RDS was calculated to be at 1:18.5 dilution.

## DISCUSSION

In this report, we demonstrate that an oral liquid of a traditional medicine, Respiratory Detox Shot (RDS), contains broad-spectrum antiviral activity, blocking the infection of SARS-CoV, SARS-CoV-2, and Influenza A viruses. We further demonstrated that RDS inhibits the early infection steps of coronaviruses. Although the detailed anti-viral mechanisms were not studied, RDS may block viral infection by directly inactivating virions or by blocking viral entry or early post-entry steps.

Anti-SARS-CoV and SARS-CoV-2 activities have also been identified in several other herbal medicines. For example, a common TCM herbal medicine *liquorice root* has been shown to contain glycyrrhizin that inhibits the replication of clinical isolates of SARS virus (*19*). In addition, another TCM for respiratory diseases, Shuanhuanglian preparation, has been shown to inhibit SARS-CoV-2 3CL protease (3CLpro) activity *in vitro* in a dose-dependent manner (*20*). Baicalin and baicalein were proposed to be the active ingredients of Shuanhuanglian for blocking 3CLpro (*20*). The active anti-viral ingredients of RDS have not been identified. However, RDS is different from baicalin and baicalein, as RDS blocks viral infection likely by directly inactivating virions or by blocking viral entry or early post entry processes, whereas baicalin and baicalein act at a later stage of the viral life cycle by blocking the activity of viral protease. Nevertheless, the *in vitro* anti-SARS-CoV-2 activity of RDS needs to be confirmed in future human clinical trails. Currently, we are conducting animal studies to determine potential *in vivo* efficacy of RDS for blocking SARS-CoV-2 infection.

## MATERIALS AND METHODS

### Cells and cell culture

HEK293T (ATCC, Manassas, VA), MDCK (ATCC, Manassas, VA), Vero E6 (ATCC, Manassas, VA), and A549(ACE2) (a gift from Virongy, Manassas, VA) were maintained in Dulbecco’s modified Eagle’s medium (DMEM) (Thermo Fisher Scientific) containing 10% heat-inactivated FBS and 1x penicillin-streptomycin (Thermo Fisher Scientific).

### Plasmid transfection and virus assembly

Lentiviral particles pseudotyped with the SARS-CoV S protein or the SARS-CoV-2 S protein were provided by Virongy LLC (Manassas, VA), or were assembled as previously described (*15*). Briefly, for the production of GFP reporter lentiviral pseudovirus, HEK293T cells were cotransfected with the vector expressing the SARS-CoV S protein or the SARS-CoV-2 S protein, pCMVΔR8.2, and pLKO.1-puro-TurboGFP. For the production of luciferase reporter lentiviral pseudovirus, HEK293T cells were co-transfected with the vector expressing the SARS-CoV S protein or the SARS-CoV-2 S protein, pCMVΔR8.2, and pLTR-Tat-IRES-Luc. Virus supernatants were collected at 48 hours post transfection, concentrated with centrifugation, and stored at -80°C. Wild-type SARS-CoV-2 virus (Isolate USA-WA1/2020) was provided by BEI Bioresources (Manassas, VA). The pHW-NA-GFP (ΔAT6) Reporter plasmid and the A/WSN/1933 H1N1-derived plasmids pHW2000-PB2, pHW2000-PB1, pHW2000-PA, pHW2000-HA, pHW2000-NP, pHW2000-NA, pHW2000-M, and pHW2000-NS were kindly provided by Dr. Feng Li. For influenza A-GFP reporter particle assembly, HEK293T cells were cotransfected with pHW2000-PB2, pHW2000-PB1, pHW2000-PA, pHW2000-HA, pHW2000-NP, pHW2000-NA, pHW2000-M, pHW2000-NS, and pHW-NA-GFP (ΔAT6). Viral supernatants were harvested at 48 hours.

### Virus infection and drug inhibition assays

RDS (a gift from Dejia Harmony, Leesburg, VA) was manufactured by Dr. Ma’s Laboratories (Burnaby, BC, Canada). All herbal ingredients in the RDS formula meets “Yin Pian” standard based on the “Chinese Pharmacopeia 2015 edition” which includes active constituents contents and limit tests of heavy metal and pesticide level. RDS is a co-decoction of herbal medicine and the final product was evaporated under vacuum conditions. SARS-CoV-2 antiserum was kindly provided by Dr. Lance A. Liotta. Arbidol-hydrochloride (Sigma) was resuspended in Dimethyl sulfoxide (Sigma). For pseudovirus infection, A549(ACE2) cells (a gift from Virongy LLC, Manassas, VA) or Vero E6 cells in 12-well plates were pre-treated with RDS for 30 minutes, infected for 4-6 hours at 37°C, and then washed and cultured in fresh medium for 48-72 hours. For the infection of Vero E6 cells, cells were also pretreated with CoV-2 Pseudovirus Infection Enhancer (CoV-2 PIE) (a gift from Virongy LLC, Manassas, VA) for another 30 minutes at 37°C following pretreatment with RDS. Cell lysates were analyzed for luciferase activity using GloMax Discover Microplate Reader (Promega). For wild-type SARS-CoV-2 infection, Vero E6 cells were pretreated with RDS for 30 minutes at 37°C, and then infected with SARS-CoV-2 (Isolate USA-WA1/2020; BEI Bioresources) at MOI of 0.05 for 1 hour inside the BSL-3 containment facility at George Mason University. Cells were washed twice with PBS and cultured for 48 hours with medium containing RDS. Virus was harvested from the supernatant and the vial titers were determined by plaque assay in Vero cell monolayers grown in 12-well plates. Briefly, serial 10-fold dilutions of each sample were prepared in complete Dulbecco’s Modified Eagle Medium (VWR) containing 1X Penicilin-Streptomycin (VWR) and supplemented with 10% FBS (Thermo Fisher Scientific). Two hundred microliters of each dilution were then adsorbed onto triplicate wells of Vero E6 cell monolayers for 1 hour. The monolayers were then overlaid with 1 to 2 ml of a mixture of 1 part 0.6% agarose (Invitrogen) and 1 part complete Eagle Minimal Essential Medium (VWR) containing 1X Penicillin-Streptomycin and supplemented with 10% FBS. At 48 hours, monolayers were fixed in 10% formaldehyde solution for 1 hour, and the overlay agar plugs were removed. To stain for plaques, 1% crystal violet dye solution containing 20% ethanol was added for 5 minutes, followed by washing with deionized water. For influenza A virus infection of MDCK cells, cells were pre-treated with RDS for 30 minutes at 37°C, and then infected with influenza A-GFP reporter virus for 6 hours. Cells were washed and cultured for 36 hours with medium containing RDS. GFP expression was quantified by flow cytometer (FACSCalibur, BD Biosciences).

### Cytotoxicity assays

Drug cytotoxicity on A549(ACE2) cells and Vero E6 cells were quantify by propidium iodide staining and flow cytometry as described (*21*). Drug toxicity on MDCK cells was quantified using Cell Proliferation Kit I (MTT) (Sigma) and the protocol suggested by the manufacturer. Briefly, MDCK cells (ATCC) were seeded into a 12 well plate at 1×10^5^ cells per well. Cells were cultured overnight, and then treated with RDS for one day, and then cultured in the medium supplemented with MTT labeling reagent (Sigma). Cells were incubated with the labeling reagent for 4 hours, followed by the addition of MTT solubilization solution. The plate was incubated overnight, and then the absorbance was measured using GloMax Discover Microplate Reader (Promega).

### Data availability

All data generated or analyzed during this study are included in this article. Reagents are available from Y.W. upon request.

## ACKNOWLEDGMENTS

The authors wish to thank Feng Li for providing influenza viral expression vectors, Lance Liotta for providing antiserum; Ted Ci, He Sun, Zhigang Gao, and Wanying Wu for discussions and advices; Kevin Carter, Mark Mamdar, Rich Keurajian, Karen Freidouni for providing RDS and herbal extracts. This work was funded by George Mason University internal grant 223741 (DeJia Harmony/Anti-SARS-CoV-2) provided by Dejia Harmony.

## AUTHOR CONTRIBUTIONS

Experiments were designed by Y.W., R.H. and L.A.H.. Manuscript was written by Y.W. and edited by L.A.H. Experiments were performed by B.H., D.Y., A.A.O., L.D.C., S.H., D.D., and Y.M.

